# SARS-CoV-2 infection and replication in human fetal and pediatric gastric organoids

**DOI:** 10.1101/2020.06.24.167049

**Authors:** Giovanni Giuseppe Giobbe, Francesco Bonfante, Elisa Zambaiti, Onelia Gagliano, Brendan C. Jones, Camilla Luni, Cecilia Laterza, Silvia Perin, Hannah T. Stuart, Matteo Pagliari, Alessio Bortolami, Eva Mazzetto, Anna Manfredi, Chiara Colantuono, Lucio Di Filippo, Alessandro Pellegata, Vivian Sze Wing Li, Simon Eaton, Nikhil Thapar, Davide Cacchiarelli, Nicola Elvassore, Paolo De Coppi

## Abstract

Coronavirus disease 2019 (COVID-19) pandemic caused by severe acute respiratory syndrome coronavirus 2 (SARS-CoV-2) infection is a global public health emergency. COVID-19 typically manifests as a respiratory illness but an increasing number of clinical reports describe gastrointestinal (GI) symptoms. This is particularly true in children in whom GI symptoms are frequent and viral shedding outlasts viral clearance from the respiratory system. By contrast, fetuses seem to be rarely affected by COVID-19, although the virus has been detected in placentas of affected women. These observations raise the question of whether the virus can infect and replicate within the stomach once ingested. Moreover, it is not yet clear whether active replication of SARS-CoV-2 is possible in the stomach of children or in fetuses at different developmental stages. Here we show the novel derivation of fetal gastric organoids from 8-21 post-conception week (PCW) fetuses, and from pediatric biopsies, to be used as an *in vitro* model for SARS-CoV-2 gastric infection. Gastric organoids recapitulate human stomach with linear increase of gastric mucin 5AC along developmental stages, and expression of gastric markers pepsinogen, somatostatin, gastrin and chromogranin A. In order to investigate SARS-CoV-2 infection with minimal perturbation and under steady-state conditions, we induced a reversed polarity in the gastric organoids (RP-GOs) in suspension. In this condition of exposed apical polarity, the virus can easily access viral receptor angiotensin-converting enzyme 2 (ACE2). The pediatric RP-GOs are fully susceptible to infection with SARS-CoV-2, where viral nucleoprotein is expressed in cells undergoing programmed cell death, while the efficiency of infection is significantly lower in fetal organoids. The RP-GOs derived from pediatric patients show sustained robust viral replication of SARS-CoV-2, compared with organoids derived from fetal stomachs. Transcriptomic analysis shows a moderate innate antiviral response and the lack of differentially expressed genes belonging to the interferon family. Collectively, we established the first expandable human gastric organoid culture across fetal developmental stages, and we support the hypothesis that fetal tissue seems to be less susceptible to SARS-CoV-2 infection, especially in early stages of development. However, the virus can efficiently infect gastric epithelium in pediatric patients, suggesting that the stomach might have an active role in fecal-oral transmission of SARS-CoV-2.

## Introduction

Severe acute respiratory syndrome coronavirus 2 is responsible for a pandemic that has proven catastrophic, due to the lack of immunity in the human population and the range of pathological features associated with infection, including severe and often life-threatening respiratory syndromes causing major health, social and economic consequences. The virus has been shown to infect respiratory epithelial cells and to spread mainly via the respiratory tract^1^. As governments and international health agencies seek effective policies to minimise infections, maintain health care delivery, and eventually ease movement restrictions, understanding the pathogenesis and the various mechanisms for transmission is of the utmost importance. It is well established that adults are more likely than children to develop symptoms upon SARS-CoV-2 infection, but little is known on the role of children in transmission of the disease. A growing body of literature suggests that replication at the level of the gastrointestinal (GI) tract not only occurs in a large proportion of confirmed cases^2^, but it also extends the overall duration of shedding, after viral clearance from the respiratory tract has occurred^3^. Interestingly, infected children have been shown to be particularly prone to develop GI symptoms which can be moderate-to-severe, leading to intensive care unit (ICU) admission and mimicking, in some cases, symptoms of appendicitis^4^. Additionally, SARS-CoV-2 was detected by means of electron microscopy in stool samples, elevated concentrations of SARS-CoV-2 RNA were detected in air samples collected in patients’ toilet areas^5^ and rectal swabs from mildly symptomatic pediatric patients persistently tested positive, even after viral clearance from the upper respiratory tract had occurred^6^. This evidence, together with the recent demonstration of a high receptor density at the level of the oral cavity and tongue^7^, raises important questions about the likelihood of fecal–oral transmission and whether therapeutic interventions to reduce gastrointestinal infection will play a role in the control of the disease. However, a recent report of 244 consecutive COVID-19 positive children from Wuhan did not find any difference in fecal nucleic acid RT-PCR between children with or without GI symptoms, suggesting that RT-PCR detection of the virus was not due to gut infection but coming instead from the respiratory tract from swallowed sputum^8^. Defining the role of the GI tract in SARS-CoV-2 infection may also help understand the risks of vertical transmission during gestation since amniotic fluid is swallowed by the fetus during gestation and viral contamination has been isolated from the placenta of an affected woman^9^. While samples from affected mothers have so far failed to prove that amniotic fluid, cord blood, and breast milk contain SARS-CoV-2^10^, very limited data are available at this stage for pregnant women with COVID-19, and even fewer data are available on intrauterine vertical transmission^11^. While severe symptoms and death have been recorded in infants as young as 10 months of age^12^, very few cases of pathology in neonates have been associated with infection^13^ and when 6 newborns from infected mothers were screened for SARS-CoV-2, they tested negative for the virus^10^. However, this limited cohort cannot exclude the possibility of infection of the fetuses.

Reliable human *in vitro* GI model systems that faithfully reproduce infection dynamics and disease mechanisms will prove key to advance our understanding of SARS-CoV-2 replication and pathology in the GI tract. Little information is available with respect to the distribution of the viral receptor angiotensin-converting enzyme 2 (ACE2) at the level of the GI tract of humans^14^. In particular, we lack fundamental information regarding which region of the GI system is the target of replication and primarily associates with the prolonged shedding of SARS-CoV-2 in both pediatric and adult patients. Organoids have attracted great attention, enabling *in vitro* disease modeling and providing an ideal tool for studying infectious pathogens, particularly of the GI tract^15^. Recent studies have demonstrated how SARS-CoV-2 can efficiently infect human intestinal enteroids^16,17^, providing evidence in support of the hypothesis that sees SARS-CoV-2 as a fecal-oral transmissible pathogen. However, it remains to be elucidated whether access to the duodenum depends on passive transport of infected oral fluids across the stomach, or on active viral replication in the gastric mucosa. Human gastric organoids derived from adult patients and induced pluripotent stem cells have proved to be instrumental for the generation of reliable *in vitro* models for the characterization of infectious agents^18,19^. Organoid derivation from human fetal organs has been shown for the intestine^20^, liver^21^ and pancreas^22^. Here, we describe the novel derivation of proliferative progenitors from human fetal stomach and their expansion *in vitro* as enterospheres. Furthermore, we provide insight into the ability of SARS-CoV-2 to infect an organoid-based model of the gastric mucosa at both fetal and pediatric ages.

This work aims to unravel the susceptibility of the stomach to SARS-CoV-2 infection through the development of an innovative expandable *in vitro* model that faithfully reproduces the gastric micro-environment. A deeper understanding of the susceptibility of the human stomach to SARS-CoV-2 infection and replication could lay the foundations for the development of therapeutic options to reduce gastrointestinal infection.

## Results

### Defined stages are present during early and late human gastric development

Organoids are organized three dimensional structures that can be grown from isolated stem cells found in adult and fetal tissues. In order to derive a novel *in vitro* gastric model of fetal origin, we firstly characterized the tissues isolated from human fetuses and compared them to gastric mucosa obtained from pediatric patients undergoing surgery (Fig. 1a). Developing stomach structures are shown in Fig. 1b from Carnegie stage (CS) 23 (corresponding to mid-week 8) to post conception week (PCW) 21. Gastric crypts start to invaginate between PCW 11 and PCW 12 and form a clearly defined crypt at around PCW 20 (Fig. 1c). We characterized the appearance of gastric markers during fetal development. Mucin 5AC positive pit mucous cells were evident at PCW 11, while pepsinogen C (marking chief cells) started to emerge at around PCW 20 (Fig. 1d). Mucin 6, a gland mucous cell marker, was constitutively expressed from early week 8 (CS 20), together with enteroendocrine cells marked by chromogranin A that were present from mid-week 8 (CS 23) (Fig. 1e). We then defined three distinct groups of gastric epithelial tissues based on gland maturity: 1) early fetal stomachs from PCW 8 to PCW 15; 2) late fetal stomachs from PCW 17 to PCW 21; 3) pediatric stomachs. Real time quantitative PCR (qPCR) was performed on gastric tissues obtained from these three groups to examine the gene expression changes of stem cell and differentiated cell markers. A significant correlation between developmental stage and mRNA expression was observed for *AXIN2*, mucin 5AC (*MUC5AC*), pepsinogen A5 (*PGA5*), with a similar trend for chromogranin A (*CHGA*) and ATPase H+/K+ transporting subunit beta (*ATP4B*) (Fig. 1f). On the other hand, expression of leucine-rich repeat-containing G-protein coupled receptor 5 (*LGR5*) and somatostatin (*SST*) were significantly higher in the late fetal stomachs.

**Figure 1.**
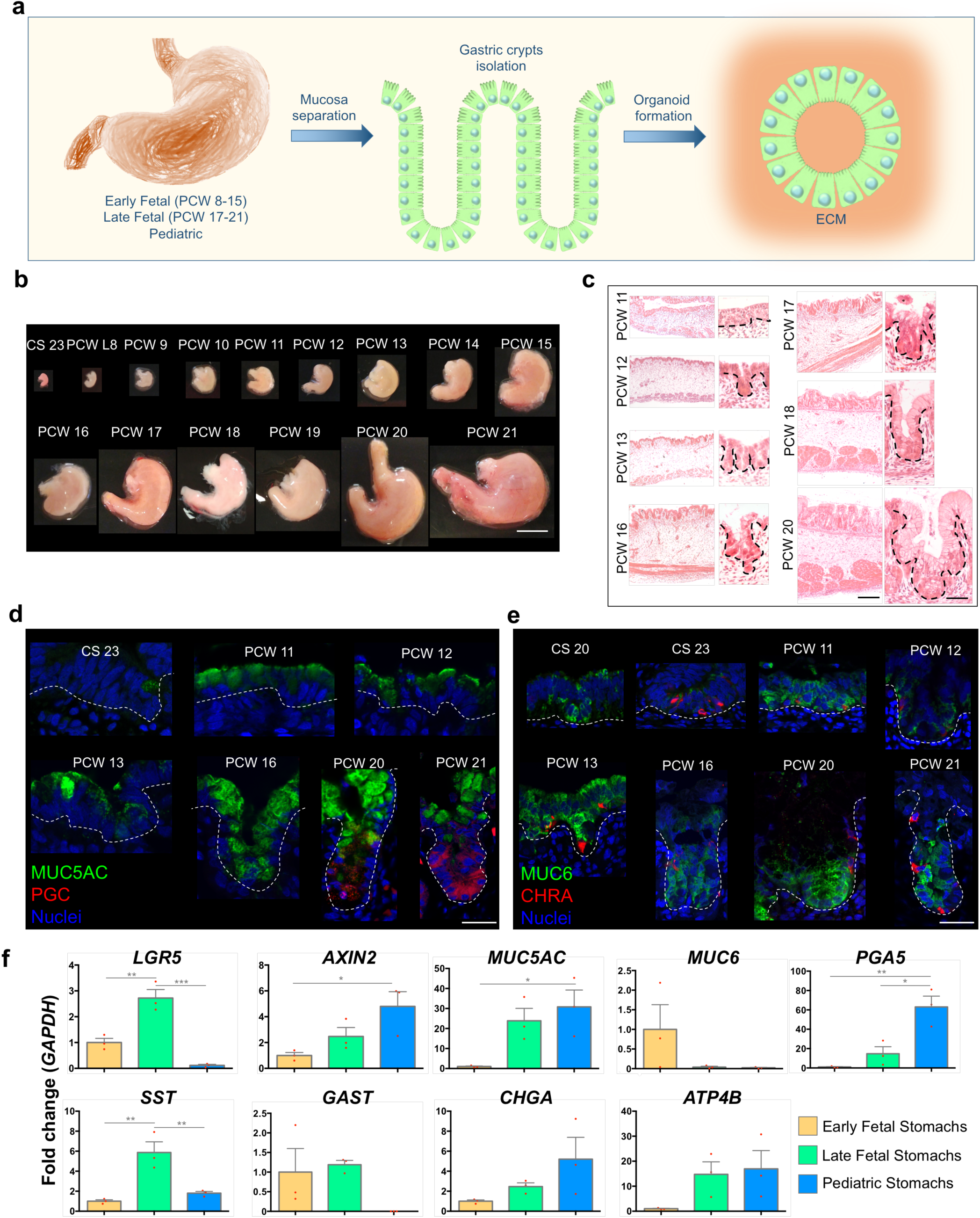
Human fetal stomach characterization. a) Schematic of the fetal and pediatric stomachs isolation, characterization and gastric organoid derivation. b) Isolated human whole stomachs from terminated pregnancies, from Carnegie stage 23 (mid-week 8) to post conception week (PCW) 21. Scale bar 1 cm. c) Hematoxylin and Eosin staining of paraffin-embedded stomach sections from PCW 11 to PCW 20. Scale bars 100 μm in lower magnification (left) and 20 μm in higher crypt magnification (right). d) Immunofluorescence panel showing mucin 5AC (MUC5AC) in green, pepsinogen C (PGC) in red and nuclei in blue (Hoechst). Scale bar 20 μm. e) Immunofluorescence panel showing mucin 6 (MUC6) in green, chromogranin A (CHRA) in red and nuclei in blue (Hoechst). Scale bar 20 μm. f) Real Time PCR analysis of early fetal, late fetal and pediatric stomachs. Relative fold change to GAPDH. Mean ± SEM (n=3). Red dots on the bar charts represent single biological replicates. Ordinary one-way ANOVA; p-value *<0.05, **<0.01, ***<0.001.

### Gastric organoids can be derived from human fetal and pediatric tissue

Following gastric tissue characterization, we efficiently extracted glandular crypts from fetal and pediatric stomachs utilizing chelating buffers and mechanical stress. To improve compatibility with subsequent clinical application of this organoid system, isolated fetal cells were expanded in a chemically defined medium, without the use of animal serum or conditioned media. Each gastric cytokine, based on previous work^19^ was screened and selectively removed from the organoids split to single cells and grown for 10 days to allow clonal organoid formation. While R-spondin 1,Wnt-3A and Noggin withdrawal led to more differentiated morphology, CHIR99021 (GSK-3 inhibitor) proved to be essential in the formation of fetal gastric organoids starting from single cells (Fig. 2a). No difference was found among fetal and pediatric organoid growth in the medium. We then performed isolation of several gastric organoid lines (Supplementary Table 1). The isolation protocol proved to be highly efficient and we obtained a biobank composed of 5 lines of early fetal stage (from CS23 to PCW 11), 6 lines of late fetal stage (from PCW 18 to PCW 21), and 4 lines of pediatric stage organoids (from 4 months-to 11 years-old). Expanding organoids were stained for the epithelial marker ezrin (EZR) and luminal polarized f-actin Fig. 2c. MUC5AC was present on the luminal side of the organoids of all stages, with a relatively lower expression in the early PCW 11 (Fig. 2c). Organoids were expanded and counted for several months, showing higher rate of expansion for earlier fetal stages (Fig. 2d). No plateau was reached in any of the curves even after several months, showing the possibility to obtain stable fetal gastric organoid lines (Supplementary Fig. 1). After weekly passaging for more than 10 weeks, we further characterized the organoid lines to evaluate genomic stability. Single nucleotide polymorphism (SNP) arrays on early fetal, late fetal and pediatric organoids showed no chromosomal duplications, no large deletions, nor other karyotype aberrations, demonstrating the organoids are genetically stable after prolonged *in vitro* culture (Fig. 2e). Real time PCR was performed on organoids grouped in early fetal (CS 23 to PCW 11), late fetal (PCW 18 to PCW 20) and pediatric. Stem cell crypt markers *LGR5* and *AXIN2* were expressed in these organoids, indicating the presence of proliferating cells. *MUC5AC* showed a pattern of increased expression along differentiation comparable to the tissue of origin in Fig. 1f. The expression patterns of *MUC6* and *SST* were also comparable to the tissue of origin. On the other hand, *CHGA* showed an inverted pattern of expression, while transcript expression of proton pump transporter *ATP4B*, responsible for gastric acid secretion, was lost in the organoid model (Fig. 2f).

**Figure 2.**
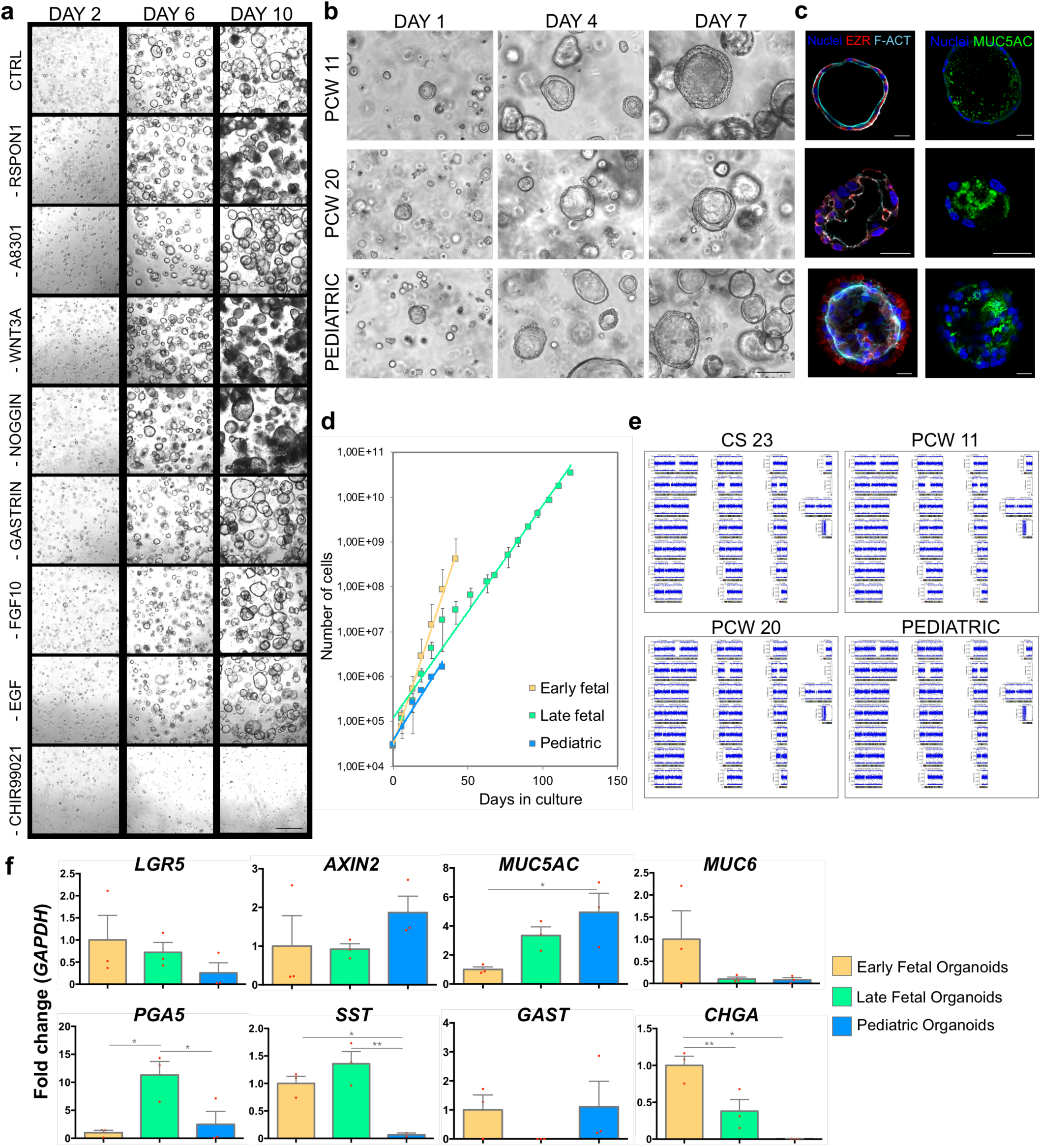
Fetal gastric organoid derivation, expansion and characterization. a) Selective withdrawal of gastric organoid cytokines from control (CTRL) complete medium, in CS 23 (mid-week 8) organoid line split at single cells at passage 7. Scale bar 400 μm. b) Bright field images of representative organoid line for each of the 3 stages, showing the formation of spherical organoids within 7 days starting from single cells. Scale bar 100 μm. c) Immunofluorescence panel showing ezrin (EZR) in red, f-actin (F-ACT) in cyan, mucin 5AC (MUC5AC) in green and nuclei in blue (Hoechst). Scale bars 25 μm. d) Cumulative cell counts during days of culture. Mean ± SD (n=4 biological replicates for early and late fetal, n=3 for pediatric organoids). e) Single nucleotide polymorphism (SNP) arrays. Representative images of the chromosome viewer and allele frequency presented for each analyzed sample. f) Real Time PCR analysis of early fetal, late fetal and pediatric gastric organoids. Relative fold change to GAPDH. Mean ± SEM (n=3). Red dots on the bar charts represent single biological replicates. Ordinary one-way ANOVA; p-value *<0.05, **<0.01.

### Gastric organoids preserve some transcriptional features of the tissues of origin

Next, we characterized the transcriptomics of gastric epithelial tissues and gastric-derived organoids, at three developmental stages. RNA-seq was performed on the three groups of early fetal, late fetal and pediatric samples. Principal component analysis (PCA) showed smaller heterogeneity in the organoid groups derived at different stages of fetal and pediatric development with respect to the primary tissues analyzed at the same stages, which may also include some heterogeneity from the surrounding cells as a result of the isolation procedure (Fig. 3a). When PCA was performed including only organoid samples, the overall variability due to the different developmental stage was comparable to that between biological replicates within the same group (Supplementary Fig. 2a). This analysis suggests that transcriptional differences related to the developmental stage of the tissue of origin could be more subtle than those captured at PCA level. We then analyzed the expression of typical gastric markers in organoids derived from tissues at different stages^23^. The only differentially expressed gene (DEG) was *MUC5AC*, which was more highly expressed in organoids from tissues at later developmental stages (Fig. 3b), confirming the qPCR results above (Fig. 2f). We did not observe processes of “intestinalization” of the organoids in culture, as *CDX2* expression was negligible (Supplementary Fig. 2c). Consistent with the qPCR results in Fig. 2f, expression of *ATP4A* and *ATP4B* proton transporters were not detectable in the RNA-seq, confirming the absence of the parietal cells in the organoids (Supplementary Fig. 2d). On the other hand, most putative genes identifying gastric crypt stem cells (*SOX9, OLFM4, PROCR, MKI67, TACSTD2*) were expressed at all developmental stages (Supplementary Fig. 2e). RNA-seq analysis on the gastric primary tissues showed a significant increase in transcript levels of the functional markers along the developmental stage (Fig. 3c), confirming that the temporal trend shown by PCA (Fig. 3a) is related to specific gastric developmental stages. When we performed hierarchical clustering analyses of the previously reported genes representing the six stomach cellular subtypes^23^, most of these genes from the analysis were not DEGs (Fig. 3d). Indeed, these six cell types are known to be all co-present at different stages of embryo development from PCW 7 to 25^23^. Furthermore, we clustered DEGs between pair of conditions to reproduce a pseudo-temporal profile between the three developmental stages considered (Fig. 3e). We highlighted in Fig. 3f the results of a pathway enrichment analysis from selected clusters that displayed gastric-related functions. Full results are reported in (Supplementary File 1).

**Figure 3:**
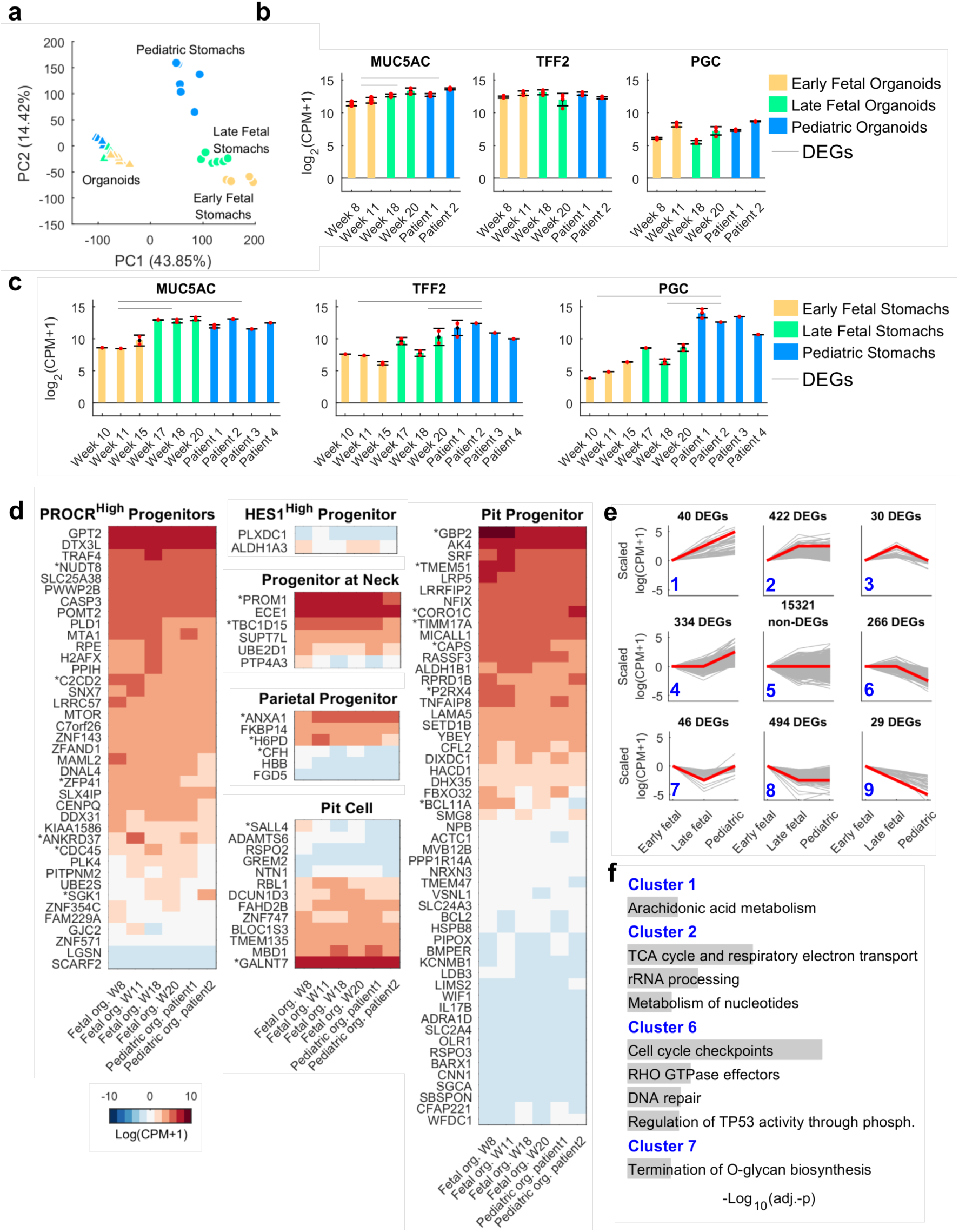
Transcriptomic characterization of gastric tissues and tissue-derived organoids from different stages of development. a) Principal Component Analysis (PCA) of RNA-sequencing samples from organoids (triangles) and fetal gastric epithelial tissues (circles) at different stages of development. Edge color identifies the biological replicates or patients within the same group of samples. Stages of fetal tissues or derived gastric organoids: early fetal (PCW 8-15), late fetal (week 17-20) and pediatric. b-c) Expression of typical gastric markers in organoids (b) and gastric epithelium (c) at different stages of development. Red dots indicate single data points. Black error bar: Mean ± SD (n = 4 for organoids biological replicates, n ≤ 3 for biological replicates tissues). CPM: counts per million. d) Results of hierarchical clustering of organoid gene expression. The selected genes were previously shown to define the six stomach cellular subtypes identified in Gao et al (2018)^23^ by single-cell RNA-seq, as indicated on top of each plot. Genes whose name is preceded by an asterisk (*) are differentially expressed genes (DEGs) between any pair of conditions. e) Pseudo-temporal profiles of gene expression according to differential expression analysis. DEGs were clustered according to a flat, increasing, or decreasing profile between pairs of time points. Data were scaled respect to the first time point. Red lines: average profile of each cluster. Gray lines: profile of each gene in the cluster. Blue digits: cluster identification number. f) Selected categories from Reactome enrichment analysis of genes in the clusters displayed in (e). Blue bars: graphical representation of the indicated values of adjusted p-value. Full results are displayed in the Supplementary Information.

### Reversed polarity in organoids facilitates expression of ACE2 on the external surface

In order to validate both fetal and pediatric gastric organoids as functional *in vitro* models of SARS-CoV-2 infection and replication, we optimized the culture condition for viral infection in a 3D system (Fig. 4a). Standard organoids of endodermal organs have a luminal polarity facing the internal portion of the structure, with an apical (inner) f-actin and zonula occludens-1 (ZO-1), and basal (external) lamina marked by β-4 integrin (β4-INT) (Fig. 4b). Such inner polarity might be an obstacle to an efficient viral infection *in vitro*, given that the apical side is luminal. In addition, Matrigel might impede efficient diffusion of the virus, thus affecting the likelihood of establishing an infection, and subsequent detection and quantification of the viral progeny released from the infected organoids. To maximize the efficiency of infection and the effective quantification of viral progeny released, we reverted the polarity of the gastric organoids^24^ to expose the apical side of the cells on the outer side. Organoids were removed from the surrounding extracellular matrix and cultured in suspension for 3 days, resulting in the exposure of the apical f-actin on the outer side, accompanied with MUC5AC secretion externally (Fig. 4c). Conversely, ZO-1 and β4-INT expression was inverted compared to standard organoids in Fig. 4b. Full 3D deconvolution images of reversed organoids are shown in Supplementary Fig. 3a.

**Figure 4.**
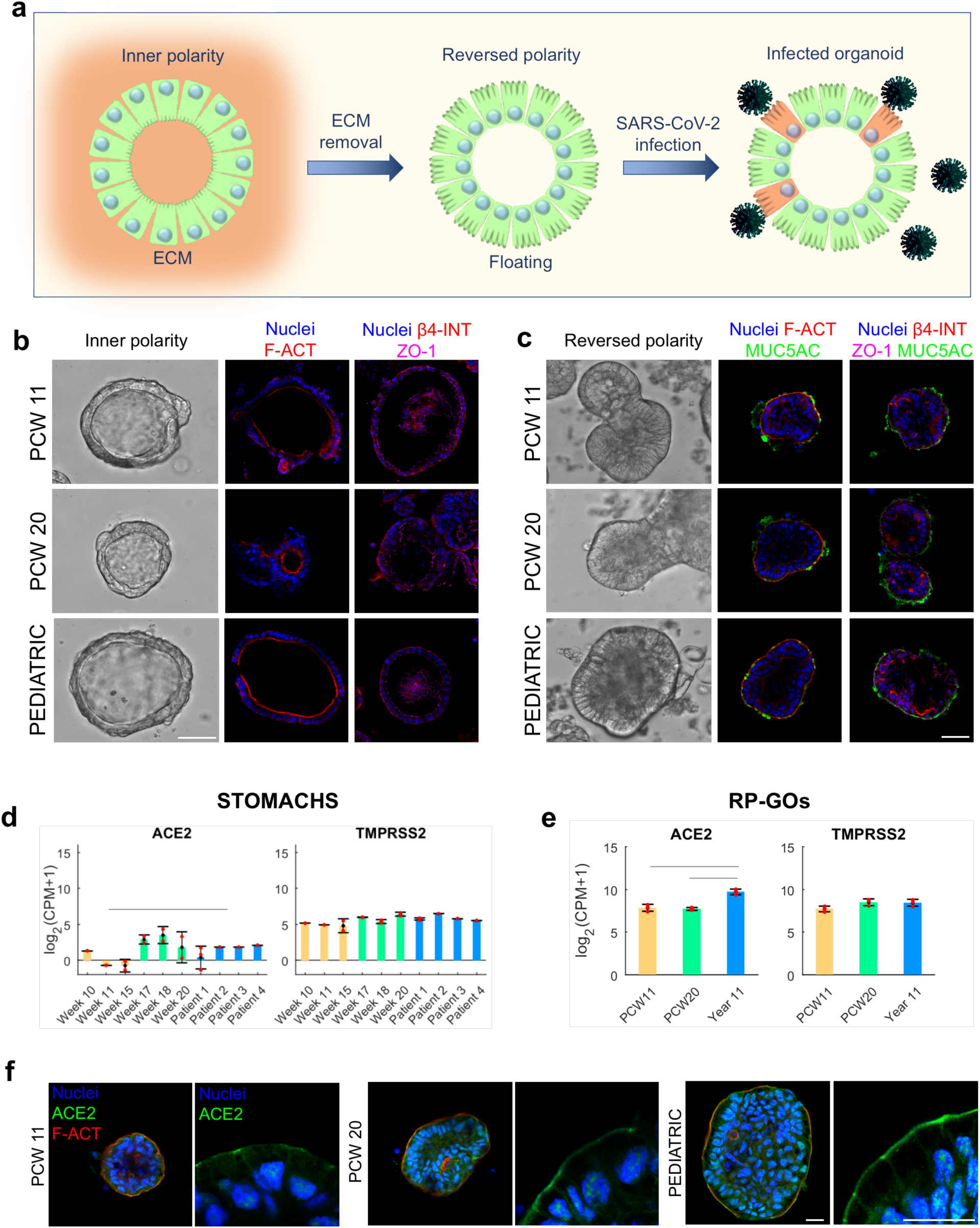
Reverse polarity organoids for efficient SARS-CoV-2 infection. a) Schematic showing the protocol for reversal of organoid polarity. ECM surrounding organoids is digested and organoids are cultured in suspension for 3 days to reverse their polarity. Once apical aspect is exposed on the outer surface, organoids are infected with SARS-CoV-2. b) Gastric organoids with normal (inner) polarity and large lumen. Immunofluorescence panel showing f-actin (F-ACT) in red, zonula occludens-1 (ZO-1) in violet, β-4 integrin (β-4 INT) in red, and nuclei in blue (Hoechst). Scale bar 50 μm. c) Gastric organoids with reversed polarity, showing an almost absent lumen. Immunofluorescence panel showing f-actin (F-ACT) in red, zonula occludens-1 (ZO-1) in violet, β-4 integrin (β-4 INT) in red, mucin 5AC (MUC5AC) in green and nuclei in blue (Hoechst). Scale bar 50 μm. d-e) SARS-CoV-2 receptors, angiotensin-converting enzyme 2 (ACE2) and transmembrane protease serine 2 (TMPRSS2) absolute RNA-seq expression in gastric tissues (left) and gastric organoids (right). Mean ± SD (n < 3 for tissues biological replicates, n = 3 for organoids). CPM: count per million. f) Immunofluorescence panel showing f-actin (F-ACT) in red, angiotensin-converting enzyme 2 (ACE2) in green and nuclei in blue (Hoechst), in fetal and pediatric gastric organoids with reversed polarity. Scale bars 20 μm.

We then analyzed the absolute expression of *ACE2* and transmembrane protease serine 2 (TMPRSS2) SARS-CoV-2 receptors in our gastric models to evaluate the SARS-CoV-2 infection potential. RNA-seq data analysis showed that expression of *ACE2* was significantly lower in early fetal stomachs compared to the pediatric ones, while late fetal samples’ higher variability prevents drawing a final conclusion. On the other hand, *TMPRSS2* mRNA expression was consistently high throughout the stomach samples (Fig. 4d). We further performed RNA sequencing on RP-GOs to evaluate the transcriptional changes. PCA analysis showed similar clustering among the different stages between RP-GOs (Supplementary Fig. 3b) and normal polarity organoids (Supplementary Fig. 2a). A comparable pattern of expression for *ACE2* and *TMPRSS2* was observed also in RP-GOs, with *ACE2* significantly higher expressed in pediatric organoids (Fig. 4e). Protein expression of ACE2 was further confirmed by immunofluorescence staining in all the RP-GOs derived at PCW11, PCW 20 and pediatric stages (Fig. 4f).

### Pediatric gastric organoids are more susceptible than fetal organoids to SARS-CoV-2 infection

To investigate the susceptibility of organoids to SARS-CoV-2 infection, we selected a SARS-CoV-2 isolate obtained from the pharyngeal swab of a 14-year-old pediatric patient. Purity of the isolate was confirmed by means of molecular testing comprising an extended panel of bacterial and viral respiratory agents. Reversed-polarity gastric organoids derived at PCW11, PCW 20 and pediatric stages and normal-polarity organoids from the same stages were infected by trained virologists in a biosafety level 3 (BSL3) laboratory. After a 2-hour infection, the organoids were cultured up to 96 hours in suspension and checked for structural integrity and viability by visual examination on a daily basis (Fig. 5a). Vero E6 cells were used as a susceptible substrate for SARS-CoV-2, to validate the infection *in vitro* (Supplementary Fig. 4a). In normal-polarity gastric organoids embedded in Matrigel, SARS-CoV-2, established an infection as demonstrated by the presence of viral double-stranded RNA (dsRNA) in the cytosolic compartment of cells (Supplementary Fig. 4b-c), nevertheless titration of the culture supernatants by FFA failed to detect viable virus at any time after infection.

**Figure 5.**
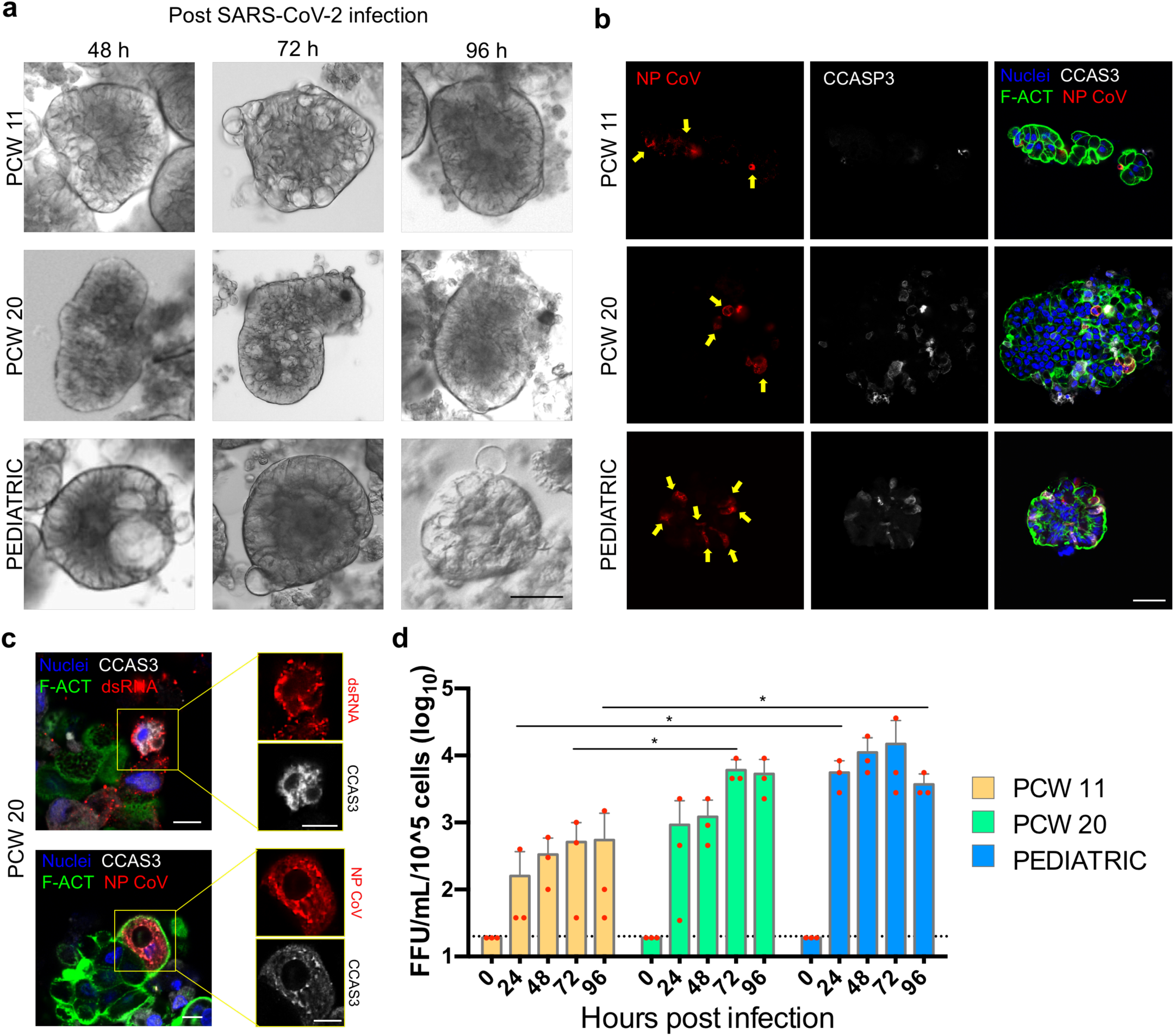
SARS-CoV-2 infection of human fetal and pediatric gastric organoids. a) Bright field images of reversed-polarity organoids infected with pediatric patient-derived SARS-CoV-2 for 2 hours, and acquired at 48, 72 and 96 h post-infection. b) Infected cells in reversed organoids fixed at 96 h post-infection. Immunofluorescence panel showing SARS-CoV-2 Nucleocapsid Antibody (NP CoV) in red, marking infected cells (yellow arrows), cleaved caspase 3 (CCAS3) in white, marking apoptotic cells, f-actin (F-ACT) in green and nuclei in blue (Hoechst). Image is representative of at least 5 similar organoid images. Scale bar 50 μm. c) Infected cell details in reversed organoids fixed at 96 h post-infection. Immunofluorescence panel showing viral double strand RNA J2 (dsRNA) in red, SARS-CoV-2 Nucleocapsid Antibody (NP CoV) in red, cleaved caspase 3 (CCAS3) in white, f-actin (F-ACT) in green and nuclei in blue (Hoechst). Scale bars 10 μm. d) Graph of SARS-CoV-2 replication in fetal and pediatric reverse-polarity organoids. Live virus yield of reverse-polarity organoids was titrated by FFA on Vero E6 cells of culture supernatants collected at 0, 24, 48, 72 and 96 hours after infection with SARS-CoV-2. The dotted line indicates the lower limit of detection. Red dots indicate single data points. Mean ± SD (N=3).

On the other hand, RP-GOs infected and cultured in suspension showed a consistent level of viral infection among organoids, as demonstrated by immunofluorescence staining of coronavirus nucleoprotein (NP CoV) and dsRNA (Fig. 5b and Supplementary Fig. 4d). Interestingly, both NP CoV and dsRNA were detected in the same cells as cleaved caspase 3 (CCAS3), meaning that SARS-CoV-2 infected gastric cells undergo programmed cell death (apoptosis), as shown in Fig. 5c. We then analyzed the level of SARS-CoV-2 replication in fetal and pediatric reverse-polarity organoids, after 0, 24, 48, 72 and 96 hours post-infection. Fig. 5d shows the yield of live virus in the culture supernatant, titrated by focus forming assay (FFA) on Vero E6 cells. These data demonstrated that the level of SARS-CoV-2 infection and the subsequent replication increased with the developmental stages of the gastric organoids, reaching in the pediatric RP-GOs peak titers of 10^4^ FFU/ml at 72 hours post-infection.

### SARS-CoV-2 infection induces a moderate antiviral state in late fetal and pediatric organoids

Next, we performed RNA-seq analysis on non-infected and infected organoids samples at each developmental stage. Interestingly, we identified significantly DEGs in samples from PCW 20 (19 DEGs) and pediatric organoids (13 DEGs), but not in PCW11 samples (Fig. 6a). All the DEGs in both developmental stages were up-regulated after the infection, among which 10 genes (*CMPK2, DDX58, DHX58, HERC5, IFI44, IFIT2, IFIT3, IRF7, MX1, RSAD2*) were in common. We further performed an analysis to find the DEGs associated with the infection irrespective of the developmental stage, identifying a further 8 DEGs (*BST2, EIF2AK2, HERC6, IFI44L, LAMP3, SLC1A3, STAT1, USP18*), for an overall number of DEGs equal to 30. Intriguingly, approximately 63% of the DEGs identified in this study as respondent to the infection were previously found to be DEGs in a literature survey of transcriptomic data on SARS-CoV, where 38 genes were identified as DEGs at the intersection of at least 9 studies (Fig. 6b). Among the common DEGs, some were first responders of the infection process, like the *DDX58* and *IFIH1* encoding the viral RNA sensors RIG-I and MDA5 respectively, and their regulators, such as *DHX58*; others more downstream players of the response, such as *OAS2* that is activated by detection of dsRNA to inhibit viral replication, *IFIT2* that inhibits the expression of viral mRNAs, and BST2 that limits viral secretion. All the 30 DEGs identified in this study, except *SLC1A3*, showed an up-regulation in response to the infection in samples from PCW 20 and the pediatric patient (Fig. 6c). *IFI44L* showed the highest fold change both in PCW 20 and in pediatric samples. This gene was previously found to be a marker of viral infection compared with to bacterial infection^25^ and more recently described as a negative modulator of innate immune responses induced after virus infections^26^. Type I, II and III interferon (IFN) transcripts were not differentially expressed between non-infected and infected RP-GOs.

**Figure 6.**
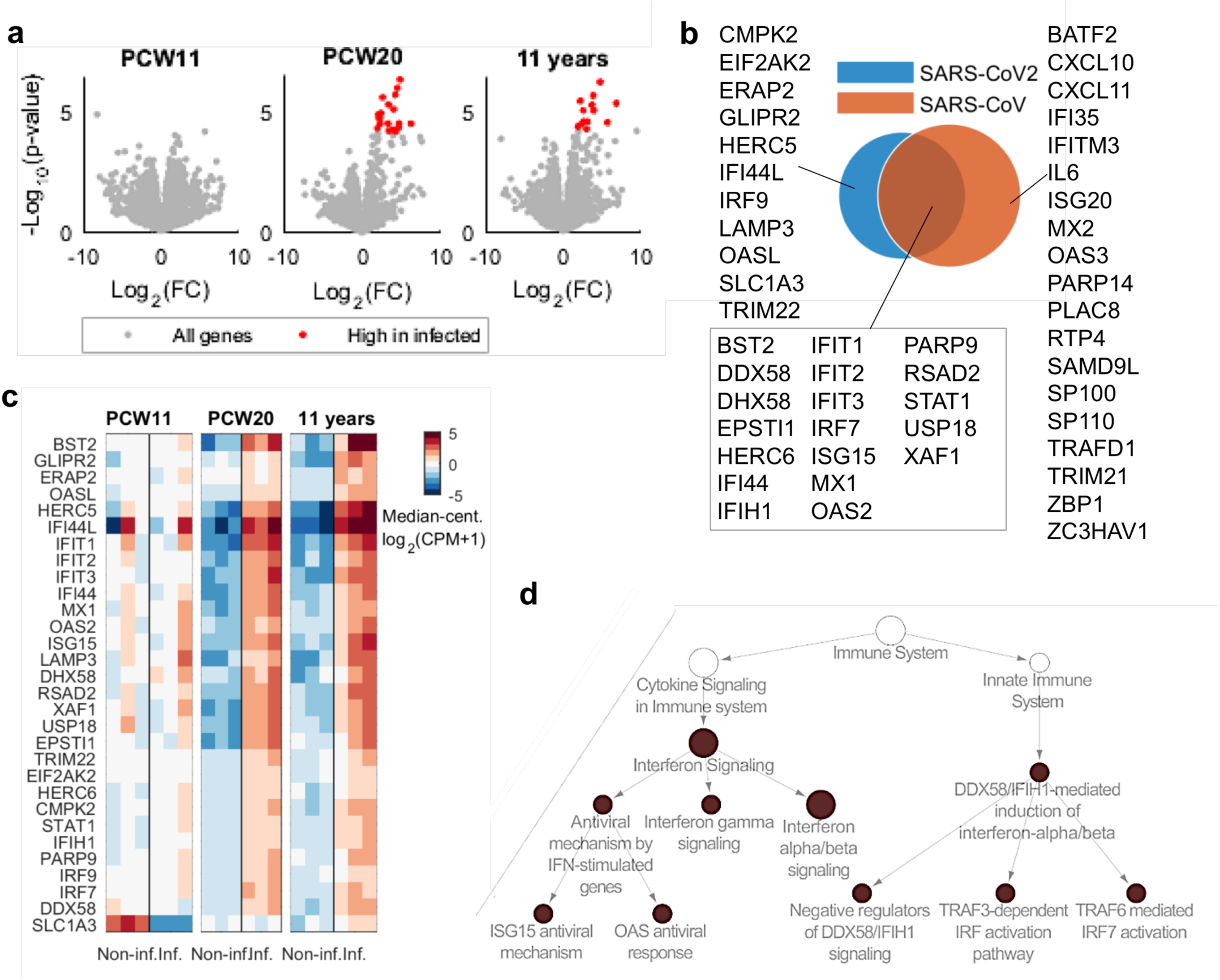
RNA-seq analysis highlights differential response to infection in organoids derived from different developmental stages. a) Volcano plots indicating DEG genes of non-infected vs. infected sample comparisons at each developmental stage (PCW 11, PCW 20, pediatric). b) Venn diagram of DEGs that responded differently to the infection in any developmental stage in this study and DEGs identified in a literature survey of transcriptomic data on SARS-CoV transfected samples^42^. c) Hierarchical clustering of DEG genes included in (b), data were median-centered for each pair of non-infected and infected conditions. d) Results from an enrichment analysis within Reactome database of DEGs highlighted in (a). Symbol size is proportional to number of genes. p-value < 10^−5^. White symbols were not enriched and were added to highlight the hierarchy between categories within Reactome structure.

We then performed an enrichment analysis within the Reactome database to understand the functional implications of the DEGs up-regulation after the infection in PCW20 and pediatric samples (Fig. 6d). Interestingly, the majority of DEGs fell within pathways associated with the innate response to viral infection, particularly those involved in the regulation of type I IFN alpha/beta by cytoplasmic pattern-recognition receptors (PRRs) such as *RIG*-I and *MDA5*, and the expression of IFN stimulated genes (*ISGs*). Moreover, to capture more subtle differences between non-infected and infected samples that do not emerge in the DEG analysis, we performed a Quantitative Set Analysis for Gene Expression (QuSAGE)^27^ within the Gene Ontology database (Supplementary File 2). Categories related to the IFN response to the infection were identified in all three sample groups (PCW11, PCW20, and pediatric), other categories were reported for completeness, but require further studies to understand their relevance.

## Discussion

Reliable *in vitro* models capable of reproducing complex *in vivo* systems are becoming increasingly important in life sciences and play a crucial role in the investigation of emerging pathogens of devastating sanitary and economic impact like SARS-CoV-2. In the context of the COVID-19 pandemic, it is still unclear how gastrointestinal virus replication might affect the clinical outcome of infection, the development of immunity and the transmission dynamics in the population. While it has been shown that SARS-CoV-2 is frequently detected in rectal samples of affected children and adults, it remains to be determined if the virus is able to produce a primary infection throughout the entire GI tract, or if its presence could be related in part to a passive transport of contaminated sputum coming from the upper respiratory tract. Most importantly, the ability of SARS-CoV-2 to persist in the GI tract after respiratory clearance, has not yet been fully elucidated in terms of viral infectivity, possibly impairing important public health and policy measures for the control of the disease. These concerns are particularly relevant in children who appear on average to suffer a less severe respiratory illness compared to adults, despite recording more prominent GI symptoms with clinical pictures mimicking appendicitis^28^, a hyperinflammatory shock syndrome (Paediatric Multisystem Inflammatory Syndrome – Temporally Associated with SARS-CoV-2, PIMS-TS)^29^, or acting as relatively asymptomatic carriers of the virus. Those risks have prompted clinical guidelines recommending the avoidance of aerosol producing procedures, including upper GI endoscopies, in children with confirmed or suspected COVID-19 for the safety of frontline clinical staff and other patients. Susceptibility of the different portions of the GI tract to SARS-CoV-2 infection has not been fully characterized and due to a paucity of autopsy reports targeting the gastric compartment^2^, the capacity of SARS-CoV-2 to infect the gastric mucosa is still unclear.

Two recent studies show that the SARS-CoV-2 receptor angiotensin converting enzyme 2 (ACE2) is highly expressed on differentiated enterocytes and that intestinal organoids derived from the small intestine can be easily infected by SARS-CoV-2^16,17^. Interestingly, intestinal organoids derived from both human and horseshoe bats are fully susceptible to SARS-CoV-2 infection and sustain robust viral replication. Although vertical transmission of SARS-CoV-2 seems to be anecdotal, it is still unclear if this lack of infection relates to the inability of the virus to migrate through the placenta^30^, to the low susceptibility of the fetal cells to infection, or simply on low viremic loads. Whereas human fetal intestinal organoids have already been reported^20^, a reliable 3D culture *in vitro* model of human gastric mucosa at different developmental stages has been challenging to achieve.

In this study, we describe successful derivation of human gastric organoids from both fetal and pediatric samples and we demonstrate that gastric cells are susceptible to SARS-CoV-2 infection. Furthermore, we describe how a reversed-polarity organoid model can help expose the apical domains, in direct contact with the surrounding microenvironment, so that pathogens can easily access surface receptors on the cells. Past studies showed laborious pathogen infection in human gastric organoids by microinjection of *Helicobacter pylori* solution into the lumen of each organoid^18,19^. Other studies showed disruption of 3D organoid organization in favor of a 2D monolayer culture to overcome inner polarity problems in *H. pylori* infection^31^. Recently, in SARS-CoV-2 infection studies, intestinal organoid 3D structures were sheared to expose the apical viral receptors and then reaggregated in ECM hydrogel droplets^16,17^. In these studies, infection of organoids upon shearing and embedding was attained, as proved by the immunofluorescent staining of viral antigens and detection of viral RNA. However, release of the infectious progeny in the culture supernatants differed greatly, ranging from positive titers of around 10^1-2^ Tissue Culture Infectious Dose 50% (TCID_50_)/mL^16^ to 10^4-5^ TCID_50_/mL^17^. Such discrepancy could depend on multiple factors, but we believe that the laborious nature of this approach might increase inter-operator and inter-laboratory variability, resulting in the generation of less-reproducible data. To overcome these complications, we decided to prevent the system perturbation and infect human gastric organoids under steady-state conditions. Taking advantage of a polarity reversion study^24^, we generated fetal and pediatric cultures of RP-GOs in suspension. In this condition of exposed apical polarity and absence of surrounding Matrigel, we could infect organoids and readily titrate the infectivity of the progeny virus, recording infection level comparable to those shown by Zhou et al.^17^, taking into account an FFU-to-TCID_50_ conversion factor of 0.28 (data not shown, from previous validation of assays). Interestingly, when we infected gastric organoids through shearing and re-embedding in Matrigel, infection was achieved but failed to detect virus in the supernatants, indicating this approach as suboptimal for our purposes.

We demonstrated that the RP-GOs are fully susceptible to SARS-CoV-2 infection, with an efficiency of replication that correlates directly with the developmental stage of origin. Quantification of gene transcripts coding for the viral receptors ACE2 and TMPRSS2 suggested that the observed levels of replication are not dependent on difference in the density of receptors, given their statistically comparable expression across the three stages. Nonetheless, variation in the protein-to-mRNA ratio across organoids of different developmental stage should be taken into account and receptivity investigated in future work. Immunofluorescence staining for the nucleocapsid indicated a clear cytosolic localization of this protein that in some cells was associated with the presence of the cleaved caspase 3, confirming the occurrence of apoptosis in the GI compartment^16^. Apoptosis in infected GI mucosal cells might account at least in part for the frequent abdominal pain, vomit and diarrhea described in COVID-19 patients^32^, in particular in pediatric populations. Apoptosis is one of the key mechanisms of cells to restrict viral infections by destruction of the cellular machinery indispensable for virus replication; on the other hand, selected viruses have evolved diverse adaptative strategies to control this phenomenon in their favor^33^. To this respect, SARS-CoV was shown to replicate *in vitro* to high titers in cells undergoing apoptosis and to low titers in cells where cytopathic effect was limited and a persistent infection was established^34^. Interestingly, induction of apoptosis for SARS-CoV was proved to be caused by a nuclear localization of the nucleocapsid protein that in turn resulted in its cleavage by caspases 6 and 3. The precise mechanism underpinning nucleocapsid cleaving, apoptosis and the replication efficiency of SARS-CoV remains unexplored. We reckon that similar mechanistic studies are of great interest to decipher the pathology of SARS-CoV-2 in the GI system and its implications on virus shedding and transmissibility.

RP-GO of late fetal and pediatric age infected with SARS-CoV-2 shared a transcriptional footprint surprisingly similar to those described for infected human small intestine enteroids^16,17^ in which type I IFN genes were either poorly expressed or undetectable, despite enterocytes and gastric cells displaying moderate levels of ISGs primarily involved in the recognition of viral RNA. Moreover, our transcriptional data are in considerable agreement with clinical and experimental profiles derived from COVID-19 patients, infected normal human bronchial epithelial cells and *in vivo* studies in ferrets^35^ that highlighted a negligible expression of genes of the IFN family but a robust expression of chemokines and ISGs. Our data provide novel evidence in support of the hypothesis that pathogenesis of COVID-19 is at least in part dependent on a reduced innate antiviral response and an unbalanced cytokine production. Nevertheless, in our model chemokines were not differentially expressed, whereas in small intestine organoids, Zhou et al^17^ reported DEGs coding both chemokine receptors and ligands. Since we conducted a bulk RNA-seq analysis and the number of infected cells in our organoids were still a minority, we speculate that many processes specific of infected cells, most likely did not reach a statistically significance level and might have gone undetected, hence imposing a cautionary approach in our interpretation of data. Interestingly, a large overlap of DEGs with previous transcriptomic studies of SARS-CoV infection was found, including the peculiar feature of a limited/absent type I IFN induction and the recruitment of a subset of cytoplasmic PRRs. Similarly to SARS-CoV, in which ORF3B and ORF6 are the main antagonists of IFN, a recent study currently under peer-revision^36^ indicates the SARS-CoV-2 ORF3B protein as a potent IFN inhibitor, supporting ours and the published transcriptomic data herein discussed.

## Conclusions

Our gastric organoid system offers a unique tool to characterize the replication of viruses and some of the associated pathological consequences of infection. This innovative model could represent an *in vitro* scalable platform for the development and testing of antiviral drug candidates targeting the GI system. A deeper understanding of the pathogenic mechanism underpinning the viral colonization of the GI system will potentially expand the available therapeutic options for the inhibition and preventing of GI infection, in an attempt to suppress viral shedding and halt spreading of the disease. The clinical importance of our findings relates to the worrisome phenomenon of prolonged shedding of SARS-CoV-2 from the GI tract and calls for further research to assess the risk of vertical transmission in infected women. Defining the susceptible age and the target anatomical sites will prove of crucial importance for the implementation of sensitive and sustainable diagnostic screening for the identification of contagious asymptomatic patients.

## Methods

### Isolation of human fetal and pediatric gastric stem cells

Human fetal stomachs were dissected from tissue obtained immediately after termination of pregnancy from 8 to 21 PCW (post conception week), in compliance with the bioethics legislation in the UK. Fetal samples were sourced via the Joint MRC/Wellcome Trust Human Developmental Biology Resource under informed ethical consent with Research Tissue Bank ethical approval (08/H0712/34+5).

Human pediatric gastric surgical biopsies were collected after informed consent, in compliance with all relevant ethical regulations for work with human participants, following the guidelines of the licenses 08ND13 and 18DS02.

Fetal stomachs and pediatric biopsies were collected in ice-cold sterile phosphate buffered solution (PBS – Sigma-Aldrich) and processed within a few hours of collection. Gastric crypt stem cells were isolated from specimens following a well-established dissociation protocol^19^. Briefly, fetal stomachs where cut open longitudinally along the lesser curvature, while ≅ 0.5 cm^2^ pediatric biopsies where processed as they were obtained. Specimens were cold-washed in a plate with chelating buffer (sterile Milli-Q water (Merck Millipore) with 5.6 mmol/L Na_2_HPO_4_, 8.0 mmol/L KH_2_PO_4_, 96.2 mmol/L NaCl, 1.6 mmol/L KCl, 43.4 mmol/L sucrose, 54.9 mmol/L D-sorbitol, 0.5 mmol/L DL-dithiothreitol, pH 7, all from Sigma-Aldrich). Mucus was removed with a glass coverslip and mucosa was stripped from muscle layer. Tissue was cut in small pieces, transferred in a 15 mL tube in new chelating buffer and pipetted repeatedly. Supernatant was discarded and 10 mL of 10 mM EDTA was added and incubated for 10 min at room temperature. EDTA was discarded and mucosa pieces were washed in ice cold PBS with Ca^2+^/Mg^2+^ (Sigma-Aldrich). Tissue was transferred to a new 10 cm plate on ice and pressure was applied on top with a sterile 3.5 cm plate, to release the crypts from the mucosa. Crypts were collected in ice-cold ADMEM+++, composed of Advanced DMEM F-12, 10 mM HEPES, 2 mM Glutamax, 1% Pen/Strep (all from Thermo Fisher Scientific). Medium with glands was filtered through a 40 μm strainer and centrifuged at 300 g for 5 min at 4°C. Supernatant was discarded and ice cold liquid Matrigel^®^ Basement Membrane Matrix Growth Factor Reduced (GFR) (Corning 354230) was added to the pellet and thoroughly resuspended. Droplets of 30 μL were aliquoted on warmed multi-well plates and incubated for 20 min at 37°C for gelation. Medium was added and changed every 3 days. For medium recipes refer to Supplementary Table 2.

### Passage of organoids

Cell were passaged every 6-8 days. To passage the organoids, Matrigel droplets were disrupted by pipetting in the well and transferred to tubes on ice. Cells were washed with 10 mL of cold basal ADMEM+++ and centrifuged at 200 g at 4°C. (First method) For single cell dissociation, supernatant was discarded, and the pellet resuspended in 1 mL of TrypLE (Thermo Fisher) and incubated for 5 min. After incubation organoids were disaggregated by pipetting, and 10 mL of ice-cold ADMEM+++ was added to dilute and inhibit TrypLE. (Second method) For standard organoid passage during expansion, the organoid pellet was resuspended in 1.5 mL of ice-cold ADMEM+++ and organoids were manually disrupted by narrowed (flamed) glass pipette pre-coated in BSA 1% in PBS, to avoid adhesion to the glass. Cells were washed, pelleted and supernatant discarded. Almost-dry pellets of disaggregated organoids (or single cells) were thoroughly resuspended in cold liquid Matrigel, aliquoted in 30 µL droplets in pre-warmed multi-well plates and incubated at 37°C for 20 min to form a gel. Rho-kinase inhibitor (Tocris) was added to single cell dissociated organoids. Medium was added and changed every 3 days.

### Polarity reversion

Fully grown gastric organoids at day 7 after single cell disaggregation were removed from surrounding extracellular matrix using a modified published protocol^24^. Matrigel was dissolved with 60 min treatment of the droplets with Cell Recovery Solution (Corning) at 4°C. Organoids were retrieved from the plates using 1% BSA-coated cut-end tips and transferred to 1% BSA-coated 15 mL tubes. Cells were extensively washed with ice-cold PBS and centrifuged at 200 g for 5 min at 4°C. Supernatant was discarded, the pellet was resuspended in complete medium and transferred to non-tissue culture treated low-adhesive multiwell plates (pre-coated in 1% BSA). Organoids were cultured in suspension for 3 days to allow reversion of polarity, before use in infection experiments.

### DNA isolation and single nucleotide polymorphism (SNP) analysis

Full grown 7 days-old organoids from passages 11 to 16 were washed from Matrigel with ice-cold PBS and DNA was extracted using DNeasy Blood & Tissue Kit (Qiagen), following manufacturer’s instructions. DNA in water was analyzed using 8 μL volume at 75 ng/μl concentration per sample. Single Nucleotide Polymorphism array Infinium Core-24 v1.1 Kit (Illumina) was used according to manufacturer’s instructions. Results were analyzed through GenomeStudio software (Illumina). We exported representative images of the chromosome viewer/B allele frequency, for each analyzed sample.

### RNA isolation

For RNA isolation from stomach tissues, i) pediatric stomach biopsies consisted of only mucosal layer from surgical samples; ii) Late fetal stomachs were cut open and mucosal layer was isolated; iii) early fetal stomachs were processed with no layer isolation, given the small size of the samples. Mucus was removed from all the samples with a glass coverslip to prevent RNA loss during the isolation protocol, and tissues washed in ice-cold PBS. Then the tissues were finely cut with a scalpel on a Petri-dish on ice and transferred to 1.5 mL tubes.

RNA was isolated from cultured organoids in Matrigel with 30 min treatment of the droplets with Cell Recovery Solution (Corning) at 4°C. Cells were then washed in ice cold PBS to remove matrix leftovers that could interfere with RNA isolation. Organoids were centrifuged at 200 g at 4°C and supernatant discarded.

Dry pellets of tissues and organoids were lysed with RLT buffer (Qiagen). RNA was isolated with RNeasy Mini Kit (Qiagen) following manufacturer’s instructions. Total RNA was quantified using the Qubit 2.0 fluorimetric Assay (Thermo Fisher Scientific).

### Real Time PCR

RNA reverse transcription was performed using the High-Capacity cDNA Reverse Transcription Kit (Thermo Fisher), according to the manufacturer’s instructions. Reverse transcription was done using the T100 thermal cycler (Bio-Rad). The qRT-PCR was performed with TaqMan gene expression assay probes (Thermo Fisher) according to the manufacturer’s instructions. The following probes (all from Thermo Fisher) were used: GAPDH (glyceraldehyde 3-phosphate dehydrogenase), LGR5 (leucine-rich repeat-containing G-protein coupled receptor 5), AXIN2 (axin-like protein), MUC5AC (mucin 5AC), MUC6 (mucin 6), PGA5 (pepsinogen A5), SST (somatostatin), GAST (gastrin), CHGA (chromogranin A), ATP4B (ATPase H+/K+ transporting subunit beta). Reactions were performed on Step One Plus Real-Time PCR System (Applied Biosystems) and results were analyzed with StepOne (Version 2.3) software (Life Technologies). GAPDH expression was used to normalize Ct values for gene expression, and data were shown as relative fold change to controls (early fetal stage), using ΔΔCt method, and presented as Mean ± SEM.

### RNA Seq and transcriptome bioinformatic analyses

For RNA-seq data of original tissues and organoids with spontaneous polarity, total RNA (100 ng) from each sample was prepared using QuantSeq 3’ mRNA-Seq Library prep kit (Lexogen GmbH) according to manufacturer’s instructions. The amplified fragmented cDNA of 300 bp in size were sequenced in single-end mode using the Nova Seq 6000 (Illumina) with a read length of 100 bp. Illumina novaSeq base call (BCL) files were converted into fastq files through bcl2fastq (version v2.20.0.422) following software guide. Sequence reads were trimmed using bbduk software (bbmap suite 37.31), following software guide, to remove adapter sequences, poly-A tails and low-quality end bases (regions with average quality below 6). Alignment was performed with STAR 2.6.0a^37^ on hg38 reference assembly obtained from cellRanger website (Ensembl 93), following online site guide. The expression levels of genes were determined with htseq-count 0.9.1 by using cellRanger pre-build genes annotations (Ensembl Assembly 93). All transcripts having <1 CPM in less than 4 samples and percentage of multimap alignment reads > 20% simultaneously were filtered out.

For RNA-seq data of non-infected and infected RP-GOs, A total of 600 pg of RNA was used as input for the synthesis of cDNA with the SMART-Seq v4 Ultra Low Input RNA Kit for Sequencing (Takara Bio USA, Mountain View, CA, USA). Manufacturer suggested protocol was followed, with minor modifications. 75 pg of DNA generated with SMART-Seq v4 Kit were used for preparation of library with NEXTERA XT DNA Library Preparation kit (Illumina Inc., San Diego, CA, USA), following suggested protocol. Libraries were sequenced in pair-end mode using a Nova Seq 6000 sequencing system on an SP, 100 cycles flow cell (Illumina Inc., San Diego, CA, USA). Illumina novaSeq base call (BCL) files were converted into fastq files through bcl2fastq (version v2.20.0.422) following software guide. Alignment was performed with STAR 2.6.0a34 on hg38 reference assembly obtained from the Gencode website (primary assembly v. 32). Transcripts estimated counts were determined with RSEM 1.3.0^38^ by using the Gencode v.32 genes annotations. All genes having <1 CPM in less than 2 replicates of the same condition were filtered out.

Differentially expressed genes (DEGs) were computed with edgeR^39^, using a mixed criterion based on p-value, after false discovery rate (FDR) correction by Benjamini-Hochberg method, lower than 0.05 and absolute log2(fold change) higher than log2(1.5). This analysis was paired between non-infected and infected samples derived from the same original sample. For RNA-seq data of organoids with spontaneous polarity, DEGs were clustered according to a flat, increasing, or decreasing profile according to the differential expression analysis between pairs of time points. Principal Component Analysis was performed by Singular Value Decomposition (SVD) on log2(CPM+1) data, after centering, using MATLAB R2017a (The MathWorks). DEGs over-representation analysis of Gene Ontology (GO) and Reactome categories was performed using ClueGO (version 2.5.4)^40^. Reactome hierarchy was visualized using ClueGO within Cytoscape^41^. Hierarchical clustering of DEGs was performed on median-centered log2(CPM+1) data in MATLAB, using Euclidean distance and complete linkage. Log-normalized expression data were analyzed by the Quantitative Set Analysis for Gene Expression (QuSAGE)^27^ Bioconductor package.

### Virus isolation and cell lines

Vero E6 cells (ATCC® CRL 1586™) were maintained in Dulbecco’s modified Eagle’s medium (DMEM, Thermo Fisher) supplemented with 10% fetal calf serum (FCS), penicillin (100 U/ml) and streptomycin (100U/ml) (all from Thermo Fisher) at 37°C in a humidified 5% CO_2_ incubator. The SARS-CoV-2 isolate was obtained from a nasopharyngeal swab collected from a 14-year-old boy in Italy. Briefly, the swab viral transport medium was filtered through a 0.22 µm filter, serially diluted and incubated onto a confluent layer of Vero E6 cells, for 5 days. To ensure purity of the viral isolate, the supernatant of the highest dilution in which cytopathic effect was visible was tested for the presence of 21 human respiratory pathogens including SARS-CoV-2, using the QIAstat-Dx Respiratory SARS-CoV-2 Panel (Qiagen). Viral stocks were produced infecting at a multiplicity of infection (MOI) of 1 Vero E6 cells cultured in DMEM supplemented with 2% FCS, penicillin (100 U/ml) and streptomycin (100U/ml) and incubating the cells for 48 hours. Supernatants were collected when 80% cells exhibited cytopathic effect and cleared by low-speed centrifugation before being stored at −80°C. All infections in this paper were performed using the third culture passage of the original isolate.

### Infection of reverse-polarity organoids in suspension

Intact organoids were embedded in two 3 µl-drops of Matrigel per well, in 24-well plates. Embedded organoids were washed once in DMEM and infected at a MOI of 0.5 by incubation with 250 µl of an expansion medium viral suspension for 2 hours. After removal of the inoculum, organoids were washed twice with a DMEM solution and 400 µl of complete medium were added to each well to maintain the culture at 37 °C with 5% CO2. Reversed-polarity organoids were infected at a MOI of 1 by incubation with 250 µl of an expansion medium viral suspension for 2 hours. After infection, organoids were washed twice in DMEM to remove unbound virus. RP-GOs were dispersed in a 400 µl expansion medium at 37 °C with 5% CO2. For all organoid cultures 50 µl of supernatant were harvested at 0, 24, 48, and 72 hours post infection. An equal volume of expansion medium replaced the sampled supernatant at each collection time. An extra sample at 96 hours post infection was collected for the RP-GOs. Samples were stored at −80°C before titration through the FFA.

### Virus titration by focus forming assay (FFA)

Supernatants of organoid cultures and aliquots of viral stocks were serially diluted and incubated on confluent monolayers of Vero E6 cells, in 96-well plates, for 1 hour. Culture medium formulation was the same used for virus propagation. After infection, the inoculum was removed and an overlay of MEM, 2% FBS, penicillin (100 U/ml) and streptomycin (100U/ml) and 0.8% carboxy methyl cellulose was added. After 27 hours, the overlay medium was removed and cells were fixed in a 4% paraformaldehyde (PFA) phosphate buffered solution (PBS), for 30 minutes at 4°C. Upon removal, cells were permeabilized by incubation with a 0.5 % Triton X-100 solution for 10 minutes. Immunostaining of infected cells was performed by incubation of the J2 anti-dsRNA monoclonal antibody (1:10,000; Scicons) for 1 hour, followed by 1-hour incubation with peroxidase-labeled goat anti-mouse antibodies (1:1000; DAKO) and a 7 min incubation with the True Blue™ (KPL) peroxidase substrate. Solution of 1% bovine serum albumin and 0.05% Tween-80 in PBS was used for the preparation of working dilutions of immuno-reagents. After each antibody incubation, cells were washed 4 times through a 5 min incubation with a 0.05% Tween-80 PBS solution. Focus forming units (FFU) were counted after acquisition of pictures at a high resolution of 4800 × 9400dpi, on a flatbed scanner.

### Staining and immunofluorescence

Human gastric tissues were fixed in 4% paraformaldehyde (PFA – Sigma-Aldrich) for 2 hours and embedded in paraffin wax, then cut at 7 μm on a microtome. Hematoxylin and Eosin (H&E) tissue slides were stained according to manufacturer’s instructions with Hematoxylin and Eosin (H&E) (Thermo Fisher).

Cultured organoids were removed from Matrigel with 30 min treatment of the droplets with Cell Recovery Solution at 4°C. RP-GOs were fixed in suspension by transferring them to 1% BSA pre-coated 1.5 mL tubes, centrifuging and resuspension in 4% PFA for 20 min in rotation. PFA was discarded and quenched with 0.1M NH_4_Cl for 1 h in rotation. Matrigel droplets (3 μL on glass slides) with embedded organoids were fixed in 2% PFA for 20 min at RT, and then washed.

Immunostaining was performed by blocking and permeabilizing the tissue slides with PBS + Triton X-100 0.1% with BSA 0.5%. Organoid whole-mounts were blocked and permeabilized with PBS + Triton X-100 0.5% with BSA 1% for 2 h at room temperature in rotation. Primary antibodies were incubated in blocking buffer for 24h at 4°C in rotation and extensively washed in PBS + Triton X-100 0.1%. Secondary antibodies were incubated overnight at 4°C in rotation and extensively washed. Slides were mounted in mounting medium, while floating organoids were moved to a glass-bottomed Petri dish and blocked with a coverslip on top. The full list of primary and secondary antibodies is presented in Supplementary table 3.

### Image acquisition

Organoids were imaged in bright field using a Zeiss Axio Observer A1. Immunofluorescence images of whole-mount staining and sections were acquired on a confocal microscope Zeiss LSM 710. Infected organoid immunofluorescence images were acquired on a Leica TCS SP5.

### Statistical analysis

Statistical analyses were performed using the following software: MATLAB (v. R2017a) for PCA, pie plot, bar plot, hierarchical clustering with proteomic and RNA-seq data. GraphPad Prism Mac (v. 6.0h) was used with all other graphs and charts.

### Financial support and acknowledgments

This research was funded by the OAK Foundation Award W1095/OCAY-14-191, the Fondazione Cariparo “Progetti di Ricerca su Covid-19”, and the Horizon 2020 grant INTENS 668294. GGG is supported by the NIHR GOSH BRC Catalyst Fellowship. BCJ is supported by the General Sir John Monash Foundation, Australia. DC is supported by Fondazione Telethon Core Grant, Armenise-Harvard Foundation Career Development Award, European Research Council (grant agreement 759154, CellKarma), and the Rita-Levi Montalcini program from MIUR and POR FESR Campania 2014-2020. NE is supported by 2018 STARS-WiC grant of University of Padova, Progetti di Eccellenza Cariparo, TWINING of University of Padova, Oak Foundation Award (Grant No. W1095/OCAY-14-191). PDC is supported by NIHR Professorship and the GOSH Children’s Charity. This research was supported by the NIHR GOSH BRC. The views expressed are those of the author(s) and not necessarily those of the NHS, the NIHR or the Department of Health. This work was supported by ShanghaiTech University. We would like to thank the Human Developmental Biology Resource (HDBR).We would like to thank Martin Sidler for his help in the collection of 3 pediatric gastric samples for analysis, and Dr Gemma Molyneux and Pei Shi Chia with their assistance with sample collections.

## Author contributions

GGG, FB, NE and PDC designed the study. GGG, NE and PDC wrote the manuscript. GGG performed the main experiments on organoid cultures. EZ characterized the fetal stomachs and derived the fetal organoids. BCJ helped in organoid characterization. FB, OG and NE designed the infection experiments. OG setup organoid culture and characterization for infection experiments. SP characterized the fetal stomachs and helped in fetal organoid derivation and expansion. FB, MP, AB and EM performed infection experiments. Ca.L performed all transcriptomic bioinformatics analyses. Ce.L and HS performed immunofluorescence analysis on infected samples. AP helped in experimental setup. DC, AM, CC and LDF performed RNA-seq and read alignments. VSWL and SE helped in experimental and manuscript discussion. NT provided HDBR license. GGG, FB, NE and PDC critically discussed the data and manuscript. All authors contributed to the final version of the manuscript.

## Data availability

The authors declare that all data supporting the findings of this study are available within the article, its Supplementary Information, attached files, and online deposited data (Gastric RNA-seq data: GSE_xxxx, it will be deposited during revision), or from the authors upon reasonable request.

## Supplementary Figures

**Supplementary Figure 1.**
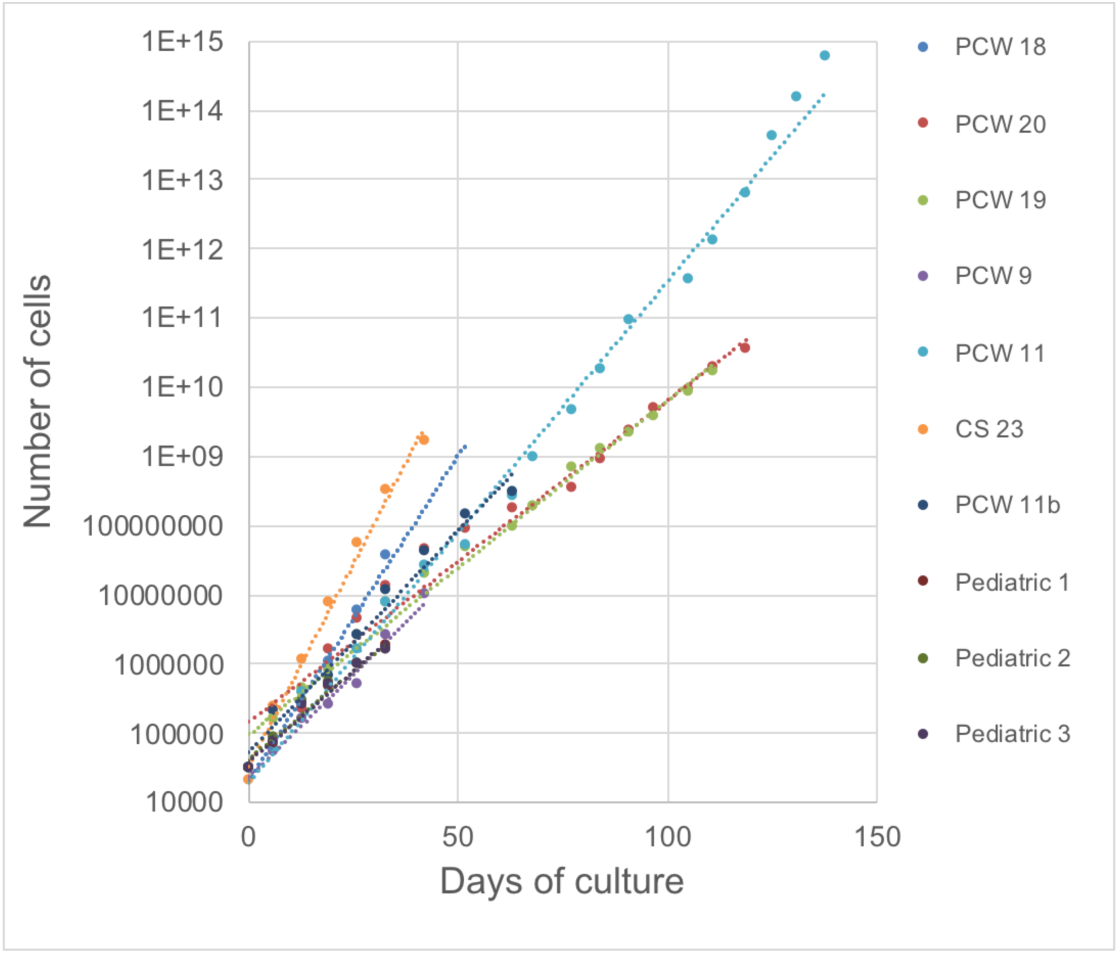
Cumulative cell counts of single organoid lines during days of culture, up to a period of 5 months.

**Supplementary Figure 2.**
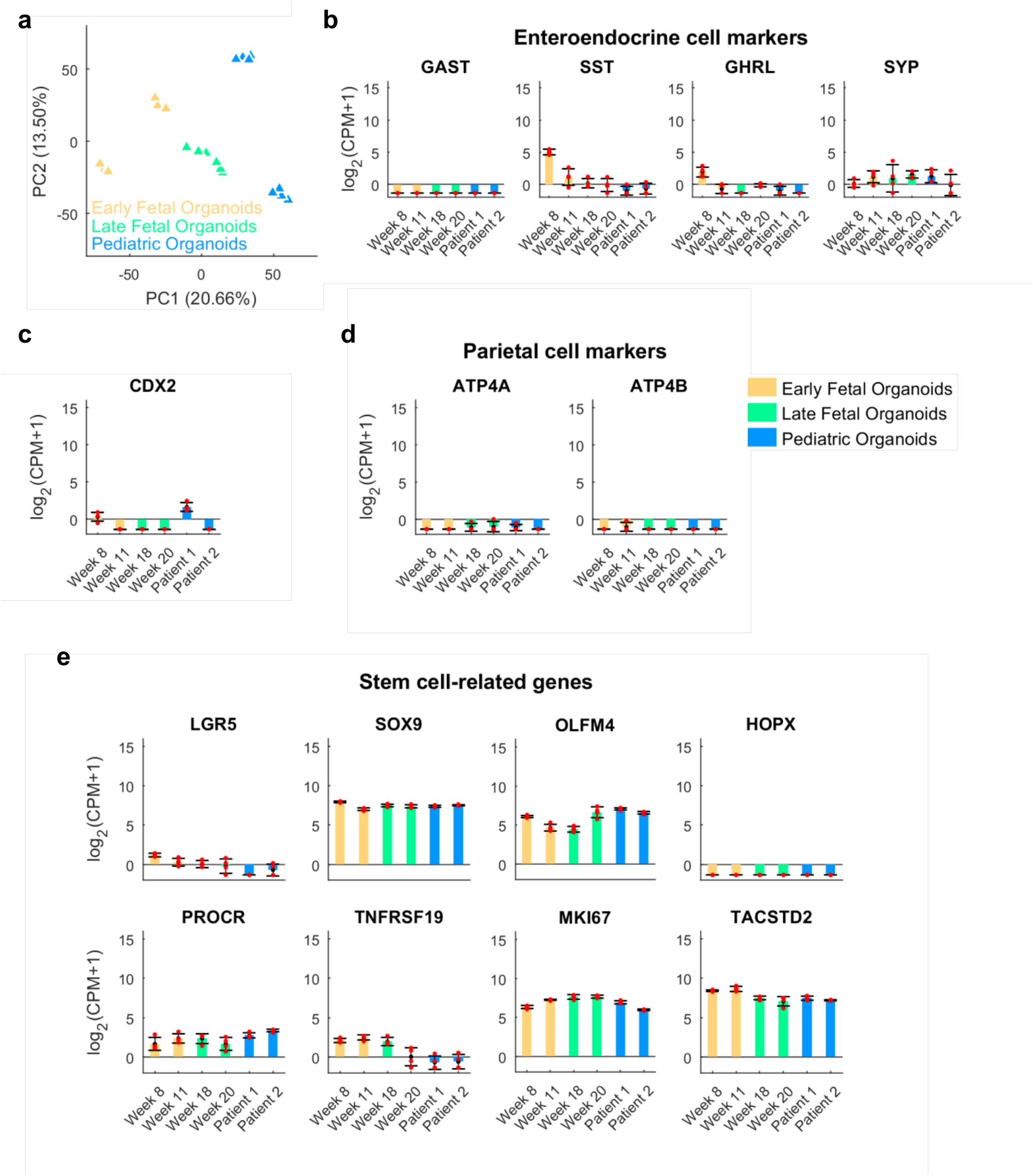
Transcriptomic characterization of organoids from different stages of development. a) Principal Component Analysis (PCA) of RNA-seq organoid samples at different stages of development as indicated. Edge color identifies the biological replicates within the same group of samples. Stages of tissue of origin for organoid derivation: early fetal (week 8-15), late fetal (week 17-20) and pediatric. b-e) Expression of selected genes in organoid samples: (b) enteroendocrine cell markers, (b) intestinal marker CDX2, (d) parietal cell markers typical of the corpus compartment, (e) putative genes identifying gastric stem cells. Red dots: single data points. Black error bar: Mean± SD (n = 4). CPM: count per million.

**Supplementary Figure 3.**
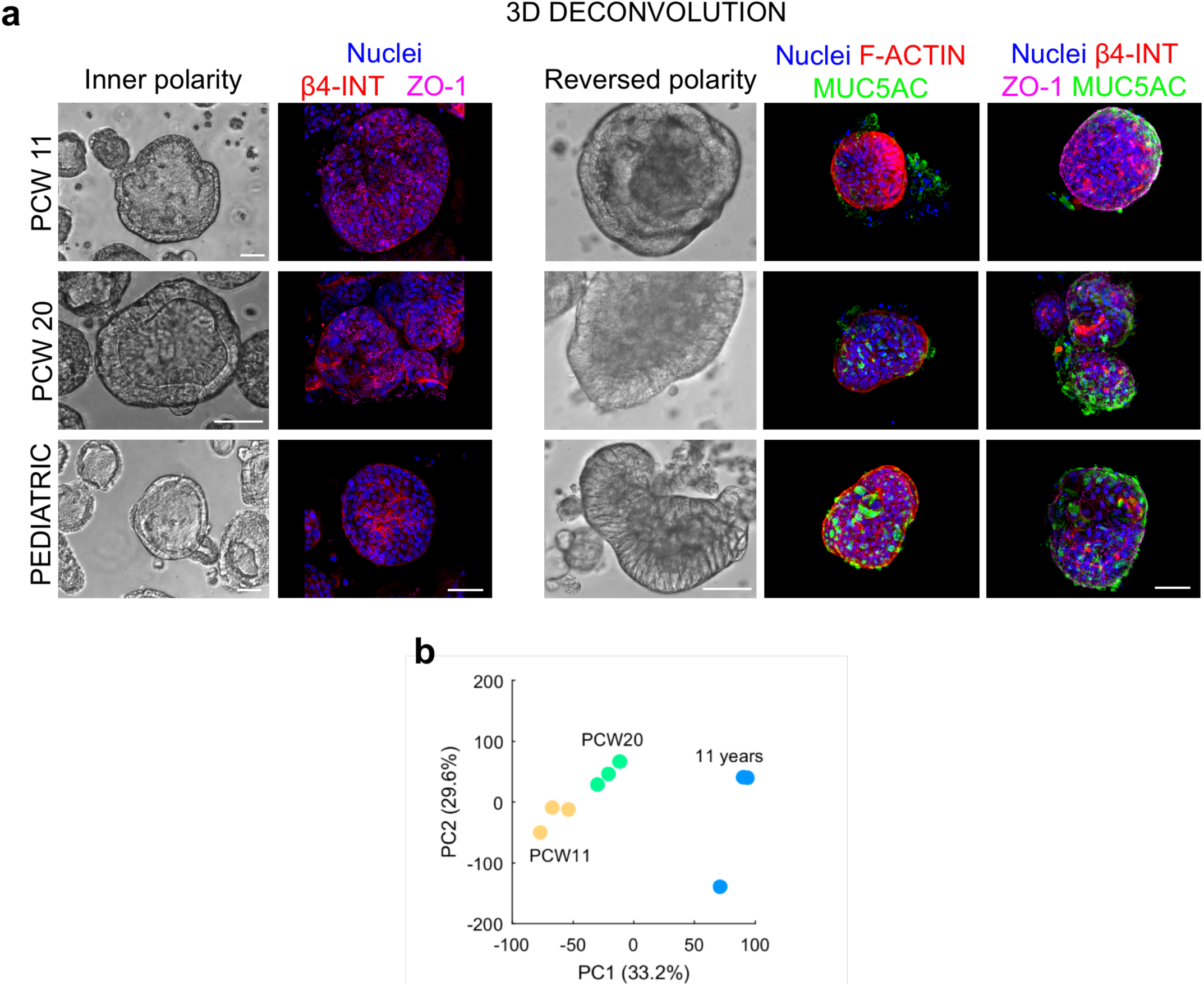
a) 3D Deconvolution confocal images. (Left) Gastric organoids with normal inner polarity and large lumen. Immunofluorescence panel zonula occludens-1 (ZO-1) in violet, β-4 integrin (β-4 INT) in red, and nuclei in blue (Hoechst). Scale bars 50 μm. (Right) Gastric organoids with reversed polarity, showing an almost absent lumen. Immunofluorescence panel showing f-actin (F-ACT) in red, zonula occludens-1 (ZO-1) in violet, β-4 integrin (β-4 INT) in red, mucin 5AC (MUC5AC) in green and nuclei in blue (Hoechst). Scale bars 50 μm. b) Principal Component Analysis (PCA) of RNA-sequencing samples from RP-GOs at different stages of development (PCW 11, PCW 20, pediatric).

**Supplementary Figure 4.**
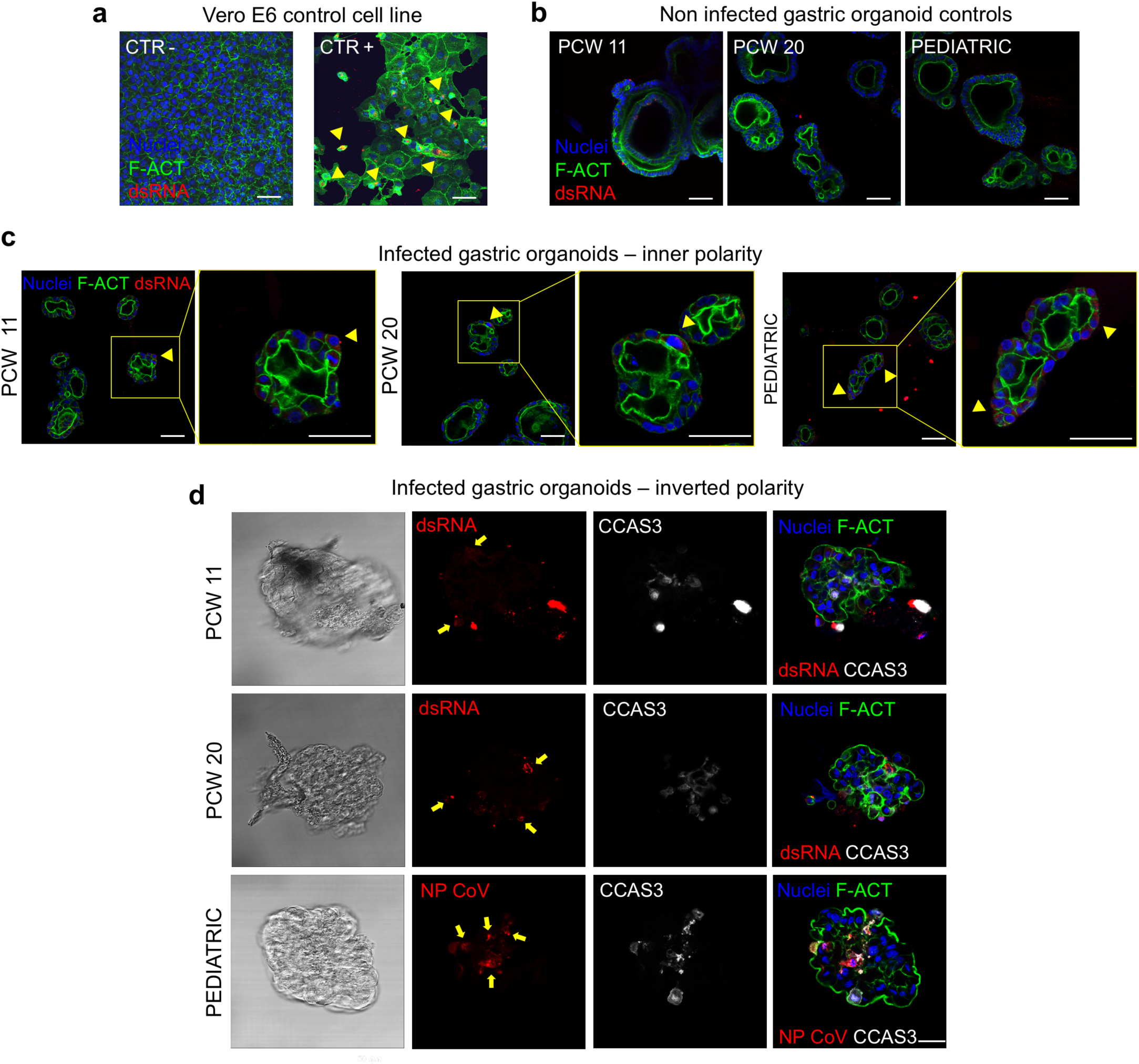
(a-c) SARS-CoV-2 staining at 64 h post-infection. a) Immunofluorescence panel on control monkey cell line Vero E6 showing viral double strand RNA J2 (dsRNA) in red, f-actin (F-ACT) in green and nuclei in blue (Hoechst). Scale bars 50 μm. b) Non-infected controls on Matrigel: whole organoids with normal polarity (right), showing absence of viral double strand RNA J2 (dsRNA) in red, f-actin (F-ACT) in green and nuclei in blue (Hoechst). Scale bars 50 μm. b) Low-efficiency infection in Matrigel whole organoids with normal polarity, showing viral double strand RNA J2 (dsRNA) in red, f-actin (F-ACT) in green and nuclei in blue (Hoechst). Scale bars 50 μm. d) SARS-CoV-2 staining at 96 h post-infection in reversed-polarity gastric organoids. Immunofluorescence panel showing viral double strand RNA J2 (dsRNA) in red, SARS-CoV-2 Nucleocapsid Antibody (NP CoV) in red, cleaved caspase 3 (CCAS3) in white, f-actin (F-ACT) in green and nuclei in blue (Hoechst). Scale bar 50 μm.

## Supplementary Tables

**Supplementary Table 1.** Gastric samples list

| Sample | Stage | Age | Organoid derivation | Use |
| --- | --- | --- | --- | --- |
| 1 | Early fetal | CS20 |  | Immuno |
| 2 | Early fetal | CS23 | Yes | Macro/Immunofluorescence/Sequencing/RTPCR/Cell count |
| 3 | Early fetal | PCW 8 (late) |  | Macro |
| 4 | Early fetal | PCW 9 | Yes | Cell count |
| 5 | Early fetal | PCW 9 |  | Macro |
| 6 | Early fetal | PCW 10 |  | Macro |
| 7 | Early fetal | PCW 10 |  | Sequencing |
| 8 | Early fetal | PCW 11 | Yes | Sequencing/RTPCR/Cell count/Immunofluorescence/Infection |
| 9 | Early fetal | PCW 11 | Yes | Expansion |
| 10 | Early fetal | PCW 11 | Yes | Cell count |
| 11 | Early fetal | PCW 11 |  | Macro/Immunofluorescence/H&E |
| 12 | Early fetal | PCW 11 |  | Macro/Immunofluorescence/H&E |
| 13 | Early fetal | PCW 11 |  | Sequencing |
| 14 | Early fetal | PCW 12 |  | Macro/Immunofluorescence/H&E |
| 15 | Early fetal | PCW 12 |  | Macro/Immunofluorescence/H&E |
| 16 | Early fetal | PCW 12 |  | Macro/Immunofluorescence/H&E |
| 17 | Early fetal | PCW 13 |  | Macro/Immunofluorescence/H&E |
| 18 | Early fetal | PCW 14 |  | Macro |
| 19 | Early fetal | PCW 15 |  | Macro |
| 20 | Early fetal | PCW 15 |  | Sequencing |
| 21 | Late Fetal | PCW 16 |  | Macro/Immunofluorescence/H&E |
| 22 | Late Fetal | PCW 16 |  | Macro/Immunofluorescence/H&E |
| 23 | Late Fetal | PCW 17 |  | Macro/H&E |
| 24 | Late Fetal | PCW 17 |  | Macro/H&E |
| 25 | Late Fetal | PCW 17 |  | Macro/H&E |
| 26 | Late Fetal | PCW 17 |  | Sequencing |
| 27 | Late Fetal | PCW 18 | Yes | Sequencing/RTPCR/Cell count |
| 28 | Late Fetal | PCW 18 |  | Macro/H&E |
| 29 | Late Fetal | PCW 18 |  | Sequencing |
| 30 | Late Fetal | PCW 19 | Yes | Cell count |
| 31 | Late Fetal | PCW 19 |  | Macro |
| 32 | Late Fetal | PCW 19 |  | Macro |
| 33 | Late Fetal | PCW 20 | Yes | Sequencing/RTPCR/Cell count/Immunofluorescence/Infection |
| 34 | Late Fetal | PCW 20 |  | Macro/Immunofluorescence/H&E |
| 35 | Late Fetal | PCW 20 | Yes | Macro/Immunofluorescence/H&E |
| 36 | Late Fetal | PCW 20 |  | Macro/Immunofluorescence/H&E |
| 37 | Late Fetal | PCW 20 |  | Sequencing |
| 38 | Late Fetal | PCW 21 | Yes | Macro/Immunofluorescence |
| 39 | Late Fetal | PCW 21 | Yes | Macro/Immunofluorescence/Cell count |
| 40 | Pediatric | 11 yo | Yes | Sequencing/RTPCR/Cell count/Immunofluorescence/Infection |
| 41 | Pediatric | 1 yo | Yes | Sequencing/RTPCR |
| 42 | Pediatric | 11 mo | Yes | Expansion |
| 43 | Pediatric | 4 mo | Yes | Expansion |
| 44 | Pediatric | 8 yo |  | Sequencing |
| 45 | Pediatric | 8 yo |  | Sequencing |
| 46 | Pediatric | 3 yo |  | Sequencing |
| 47 | Pediatric | 8 yo |  | Sequencing |
| 48 | Pediatric | 4 yo |  | Sequencing |
| 49 | Pediatric | 13 yo |  | Sequencing |

**Supplementary Table 2.** Human fetal and pediatric gastric organoid medium

| Component | Stock conc. | Final conc. |
| --- | --- | --- |
| Advanced DMEM F-12 (Thermo 12634) | - | To volume |
| HEPES (Thermo 15630080) | 1 M | 10 mM |
| Glutamax (Thermo 35050061) | 100 X | 2 mM |
| Pen/Strep (Thermo 15140122) | 100 % | 1 % |
| Primocin (Thermo Fisher NC9392943) | 50 mg/mL | 2 µL/mL |
| B-27 supplement minus vitamin A (Thermo 12587010) | 50 X | 1 X |
| n-acetylcysteine (Sigma A9165) | 500 mM | 1.25 mM |
| Wnt-3A (Peprotech 315-20) | 50 µg/mL | 100 ng/mL |
| R-spondin 1 (Peprotech 120-38) | 100 µg/mL | 500 ng/mL |
| Noggin (Peprotech 120-10C) | 100 µg/mL | 100 ng/mL |
| EGF (Thermo Fisher PMG8043) | 500 µg/mL | 50 ng/mL |
| Gastrin (Sigma G9020) | 100 µM | 10 nM |
| GSK-3 inhibitor (CHIR 99021) (Tocris 4423) | 3 mM | 3 µM |
| TGFb inhibitor (A83-01) (Sigma SML0788) | 500 µM | 5 µM |
| FGF10 (Peprotech 100-26) | 100 µg/mL | 200 ng/mL |
| ROCK inhibitor Y-27632 (Tocris 1254) | 10 mg/mL | 10 µg/mL |

**Supplementary Table 3.** Antibody list

| Antibody | Code | Brand | Host | Dilution |
| --- | --- | --- | --- | --- |
| Chromogranin A (CHGA) | ab15160 | Abcam | Rabbit | 1:50 |
| Mucin 5AC (MUC5AC) | ma5-12178 | Invitrogen | Mouse | 1:100 |
| Mucin 6 (MUC6) | ab216017 | Abcam | Mouse | 1:100 |
| Pepsinogen (PGC) | HPA031717 | Atlas | Rabbit | 1:100 |
| Ezrin (EZR) | PA5-29358 | Thermo Fisher | Rabbit | 1:200 |
| Zonula occludens-1 (ZO-1) | 40-2200 | Life Technologies | Rabbit | 1:200 |
| Integrin beta-4 (INTB4) | ab110167 | Abcam | Rat | 1:200 |
| Cleaved Caspase-3 (CCAS3) | 9661 | Cell Signaling | Rabbit | 1:400 |
| Angiotensin-converting enzyme 2 (ACE2) | MAB933 | R&D Systems | Mouse | 1:100 |
| Double Strand RNA (dsRNA J2) | J2 | Scicons | Mouse | 1:200 |
| SARS-CoV/SARS-CoV-2 Nucleocapsid (NP CoV ) | 40143-MM05 | Sino Biological | Mouse | 1:200 |
| anti-Mouse 647 (secondary AB) | A32788 | Thermo Fisher | Donkey | 1:300 |
| anti-Rabbit 647 (secondary AB) | A32795 | Thermo Fisher | Donkey | 1:300 |
| anti-Rat 568 (secondary AB) | A11007 | Thermo Fisher | Goat | 1:300 |
| anti-Rabbit 594 (secondary AB) | A11012 | Thermo Fisher | Donkey | 1:300 |
| anti-Rabbit 568 (secondary AB) | A11011 | Thermo Fisher | Goat | 1:300 |
| anti-Rabbit 488 (secondary AB) | A11008 | Thermo Fisher | Goat | 1:300 |
| anti-Mouse 488 (secondary AB) | A11001 | Thermo Fisher | Goat | 1:300 |
| anti-Mouse 568 (secondary AB) | A10037 | Thermo Fisher | Goat | 1:300 |
| Phalloidin (f-actin) | A22287 | Thermo Fisher | --- | 1:100 |
| Phalloidin (f-actin) | A12379 | Thermo Fisher | --- | 1:100 |
| Hoechst 33342 (nuclear staining) | H1399 | Thermo Fisher | --- | 10 µg/mL |

